## Supplementary Figures for "Communication between DNA polymerases and Replication Protein A within the archaeal replisome"

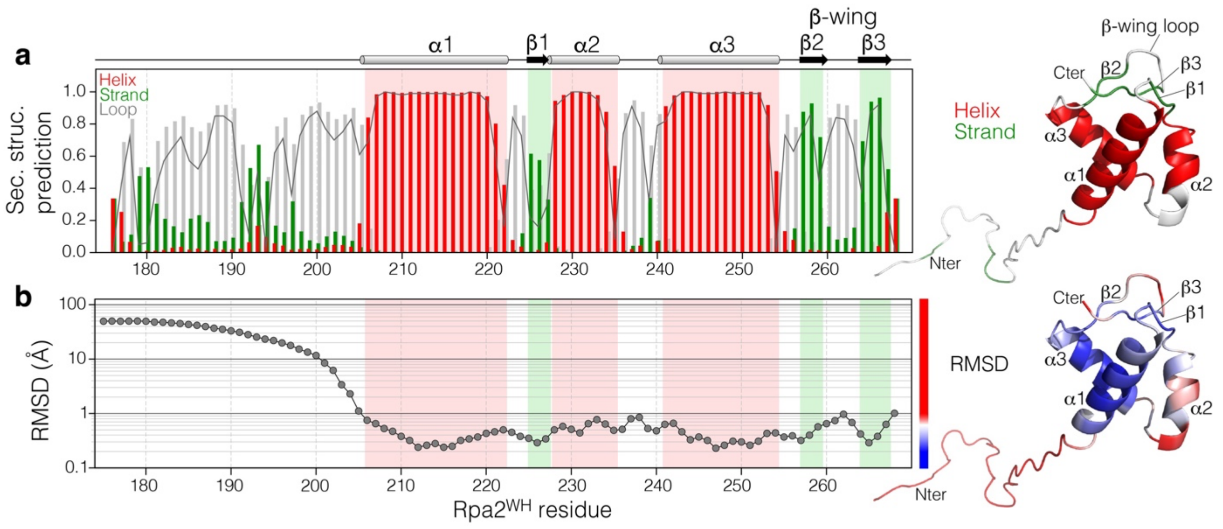

**Supplementary Figure 1**: **(a)** Secondary structure prediction of Rpa2$^{WH}$ from TALOS-N software based on HN, Hα, Cα, Cβ, CO and N secondary shifts, with predicted helices in red, strands in green and loop/coil regions in grey. The confidence is depicted by the dark grey line. Secondary structure elements inferred from these predictions are indicated at the top. **(b)** Backbone RMSD of the NMR structure ensemble. On the right of each graph, mappings of the secondary structure elements and RMSD are shown on the representative structure of Rpa2$^{WH}$, with the color coding as indicated.

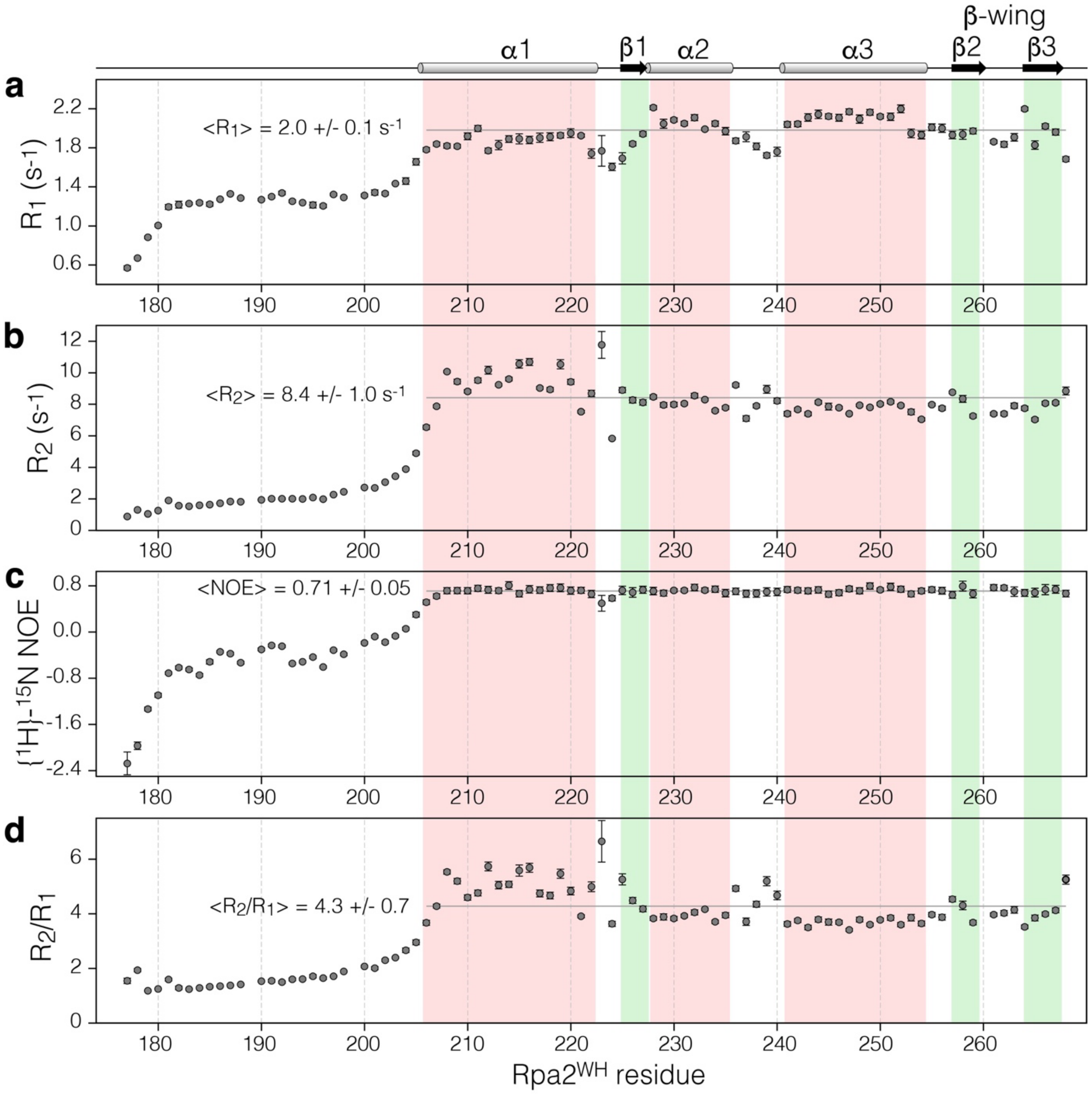

**Supplementary Figure 2**: **$^{15}$N relaxation parameters of Rpa2$^{WH}$ at 35°C, 600 MHz**. $R_1$ **(a)**, $R_2$ **(b)**, {$^1$H}-$^{15}$N NOE **(c)** and $R_2/R_1$ ratio **(d)**. Averages and standard deviations are indicated. Secondary structure elements are displayed at the top; helices and strands are highlighted in red and green, respectively.

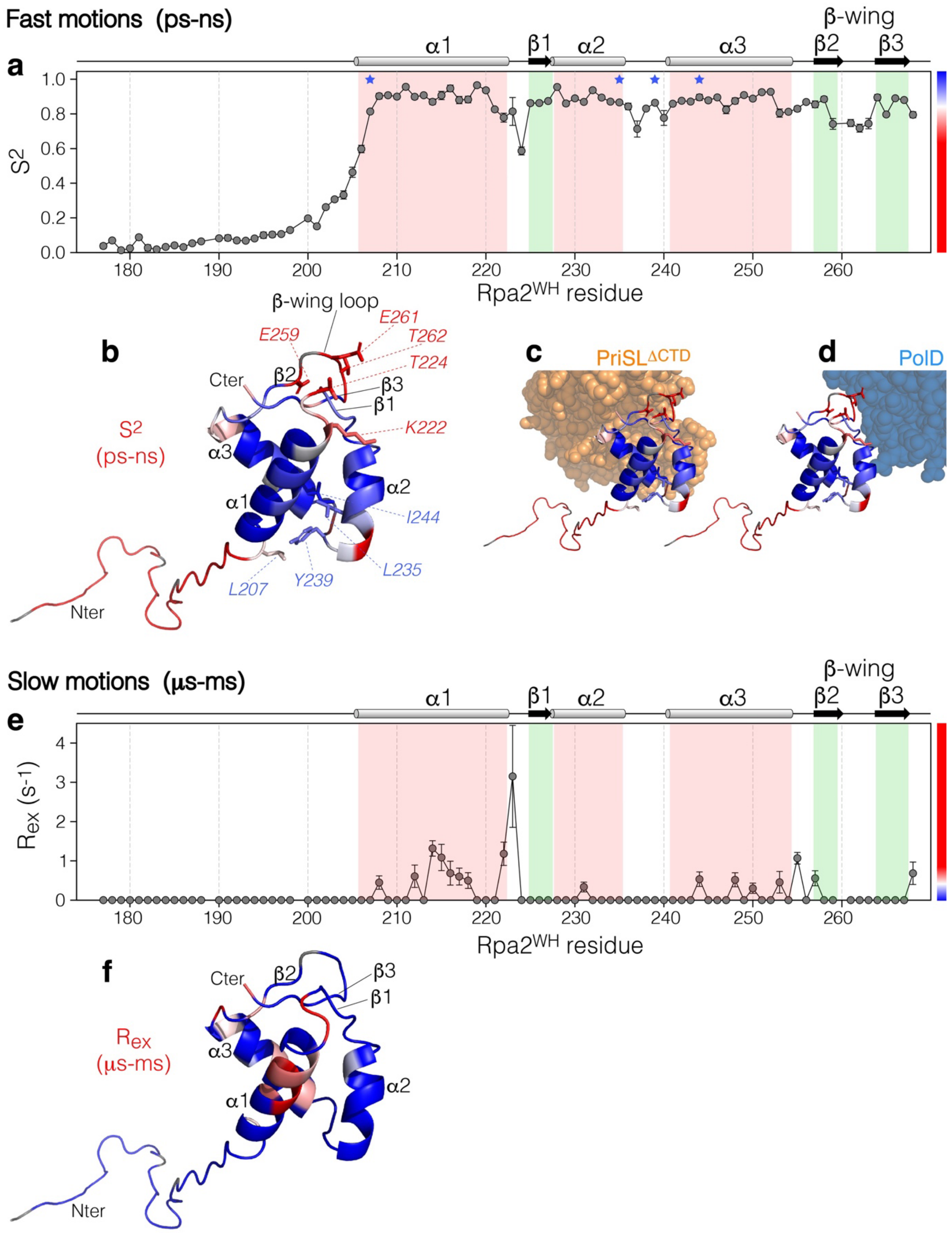

**Supplementary Figure 3: Backbone dynamics parameters of Rpa2$^{WH}$**, extracted from $^{15}N$ relaxation data using the Lipari-Szabo model-free approach with an anisotropic global reorientation model: **(a)** Order parameter $S^2$ probing motions on the ps-ns timescale. Secondary structure elements are indicated at the top and helices and strands are highlighted in red and green, respectively. **(b)** Fast motions ($S^2$) mapped on the representative structure of Rpa2$^{WH}$, with the color coding as indicated on **(a)**, i.e., from blue to red for increasing ps-ns dynamics.

**(c, d)** Close up view of the binding interface of $Rpa2^{WH}$ with $PriSL^{\Delta CTD}$ (orange spheres) and PolD (blue spheres). **(e)** Exchange contribution $R_{ex}$ probing motions on the µs-ms timescale. **(f)** Slow motions ($R_{ex}$) mapped on the representative structure of $Rpa2^{WH}$, with the color coding as indicated on **(e)**, i.e., from blue to red for increasing µs-ms dynamics. **Rationalization of the dynamics:** The N-terminal region up to residue 205 is highly flexible. The dynamics of the folded WH domain can be summarized as follows: on the N-terminus side of the domain, the loop between α2/α3 (236-240) has restrained dynamics on the ps-ns timescale due to the stabilization induced by the packing of Y239 into the hydrophobic pocket defined by L207, L235 and I244 (blue asterisks on **(a)** and blue labels on **(b)**), anchoring the N-terminus of α1 to the C-terminus of α2 and to α3 directly after the long flexible N-terminus tail. Opposite to the N-terminus, the region encompassing the loop between α1 and β1 (K222-T224) and the β-wing loop (259-263) define a flexible hotspot (red labels on **(b)**, with $S^2 < 0.75$. Interestingly, this flexible region of $Rpa2^{WH}$ lies against $PriSL^{\Delta CTD}$ **(c)** and PolD **(d)** and is involved in many electrostatic interactions with each partner as observed in the XR and cryo-EM structures of the respective complexes. It is likely that the dynamic nature of this hotspot around the β-wing helps driving fast binding to the cognate polymerases and that this region might be stabilized/rigidified upon complex formation. Conformational exchange in the µs-ms timescale is detected for residues of helices α1 and α3 pointing towards the core of the domain ($R_{ex} \sim 1$ $s^{-1}$, **(e,f)**). This phenomenon could arise from the propagation of the fast dynamics of the N-terminal tail, slowing down towards the core.

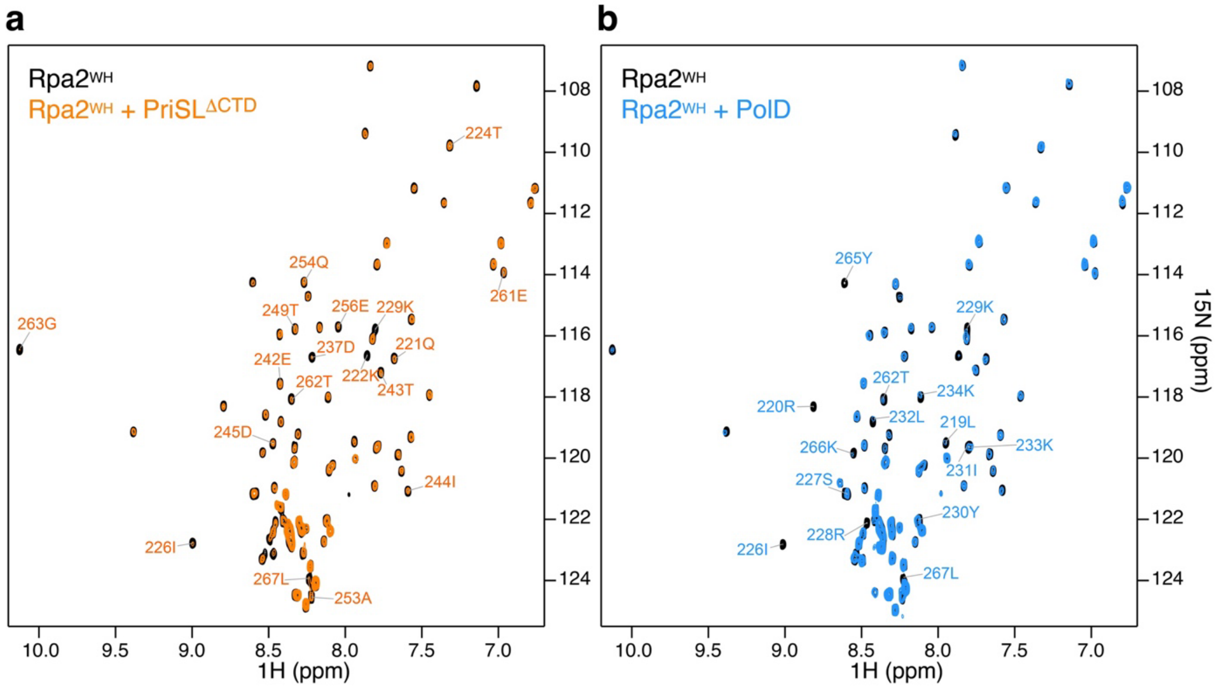

**Supplementary Figure 4**: Superimposed $^{1}$H-$^{15}$N HSQC spectra recorded on 100 µM $^{15}$N-labeled Rpa2$^{WH}$ (**a, b,** black), with 0.5 equivalents of unlabeled PriSL$^{\Delta CTD}$ (**a**, orange) or with 0.1 equivalent of unlabeled PolD (**b**, blue). Assignment is indicated for the most perturbed Rpa2$^{WH}$ signals by PriSL$^{\Delta CTD}$ ($I_{cplx}/I_{free}$ <0.29) or by PolD ($I_{cplx}/I_{free}$ <0.26). These thresholds were used to delineate the respective binding surfaces to the polymerases on **Fig. 3 k, l**.

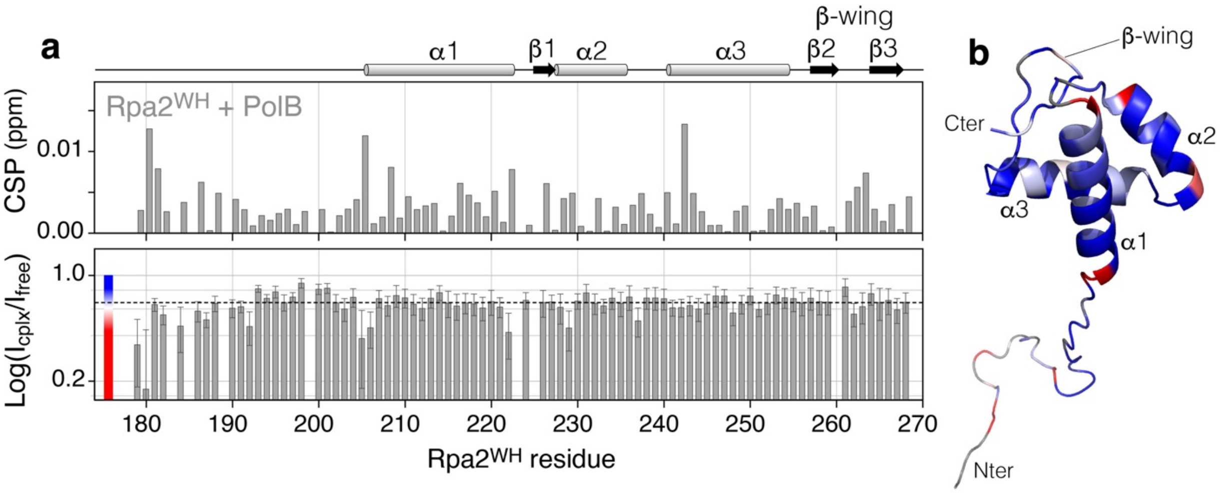

**Supplementary Figure 5**: **(a)** NMR chemical shift perturbations (CSP) and peak intensity ratios $I_{cplx}/I_{free}$ ($log_{10}$ scale) on Rpa2$^{WH}$ induced by PolB. The average $<I_{cplx}/I_{free}>$ in the folded domain (0.66) is indicated by the dotted line. $^{1}H$-$^{15}N$ HSQC spectra were recorded on 50 μM $^{15}N$-labeled Rpa2$^{WH}$ with and without an equimolar amount of unlabeled PolB. Secondary structure elements are indicated at the top. **(b)** Mapping of the intensity ratio $I_{cplx}/I_{free}$ on Rpa2$^{WH}$ with the color coding from blue (no attenuation) to red (large attenuation) as indicated. **Rationalization of PolB binding:** Despite an equimolar addition of PolB, the average $I_{cplx}/I_{free}$ ratio (0.66 ± 0.07) in the folded domain is much higher than for the complexes with only 0.5 equivalents of PriSL$^{\Delta CTD}$ (0.32 ± 0.05) or with 0.1 equivalent of PolD (0.31 ± 0.09). In addition, no significant and localized CSP or peak intensity change is detected outside of error bars on Rpa2$^{WH}$ upon addition of PolB. Taken together, these observations indicate that, unlike with the PriSL$^{\Delta CTD}$ and PolD polymerases, no strong and specific binding of the Rpa2$^{WH}$ occurs with PolB.

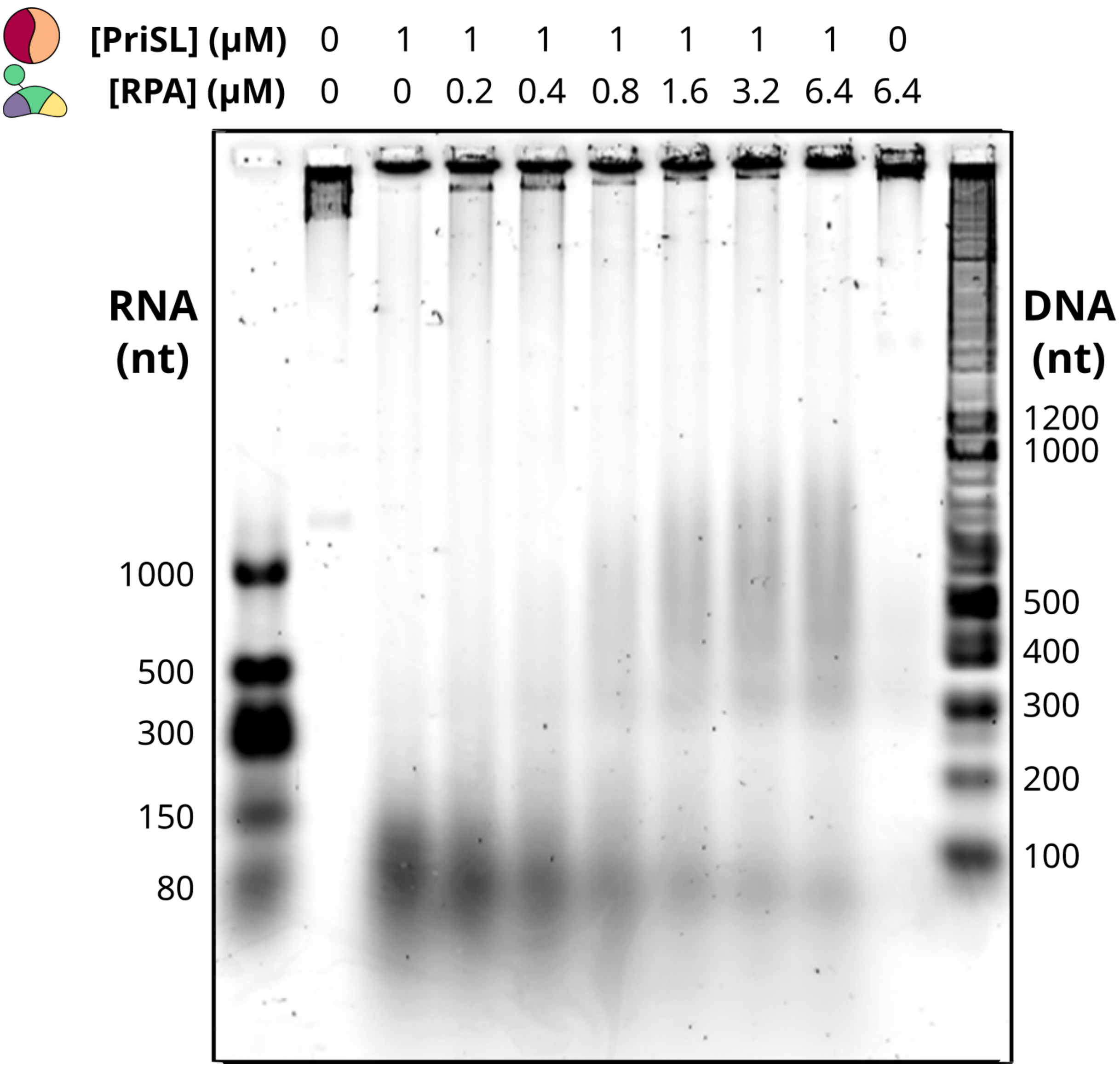

**Supplementary Figure 6**: **Impact of RPA binding on PriSL priming activity.** PriSL (1 μM) was incubated with increasing amounts of RPA (lanes 4-9). Lane 2 is the negative control without proteins. Lane 3 contains 1 μM of PriSL. Lane 10 contains 6.4 μM WT-RPA. Lanes 1 and 11 contain oligonucleotide 1 kb Plus DNA Ladder and Low Range ssRNA Ladder, respectively.

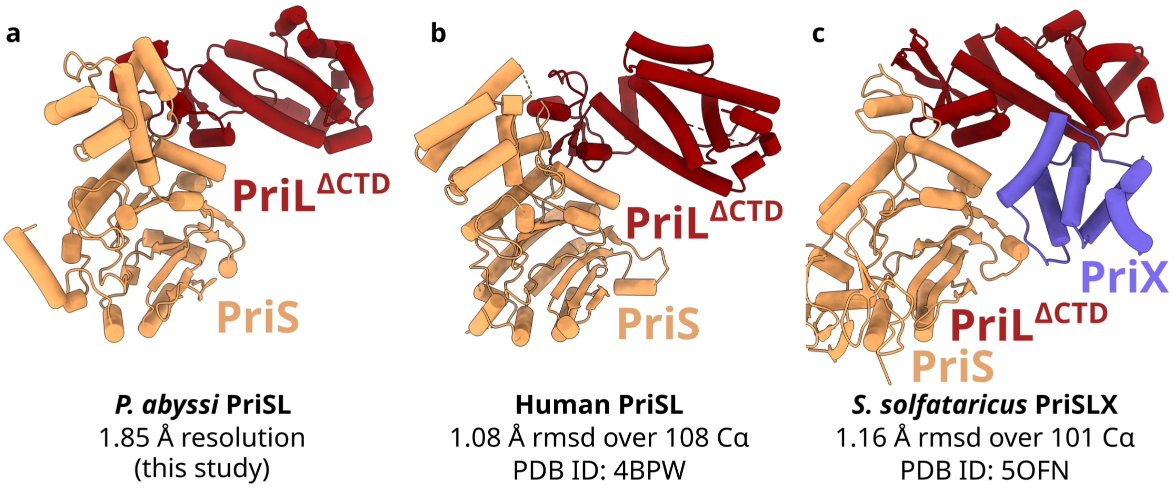

**Supplementary Figure 7**: **Crystal structure of the heterodimeric euryarchaeal DNA primase. (a)** Model of *Pyrococcus abyssi* PriS-PriL$^{\Delta CTD}$ crystal structure at 1.85 Å resolution (PabPriSL$^{\Delta CTD}$). (**b**) Alignment of human primase (PDB ID 4BPW) to PabPriSL$^{\Delta CTD}$. (c) Alignment of *Saccharolobus solfataricus* PriSLX (PDB ID 5OFN) to PabPriSL$^{\Delta CTD}$.

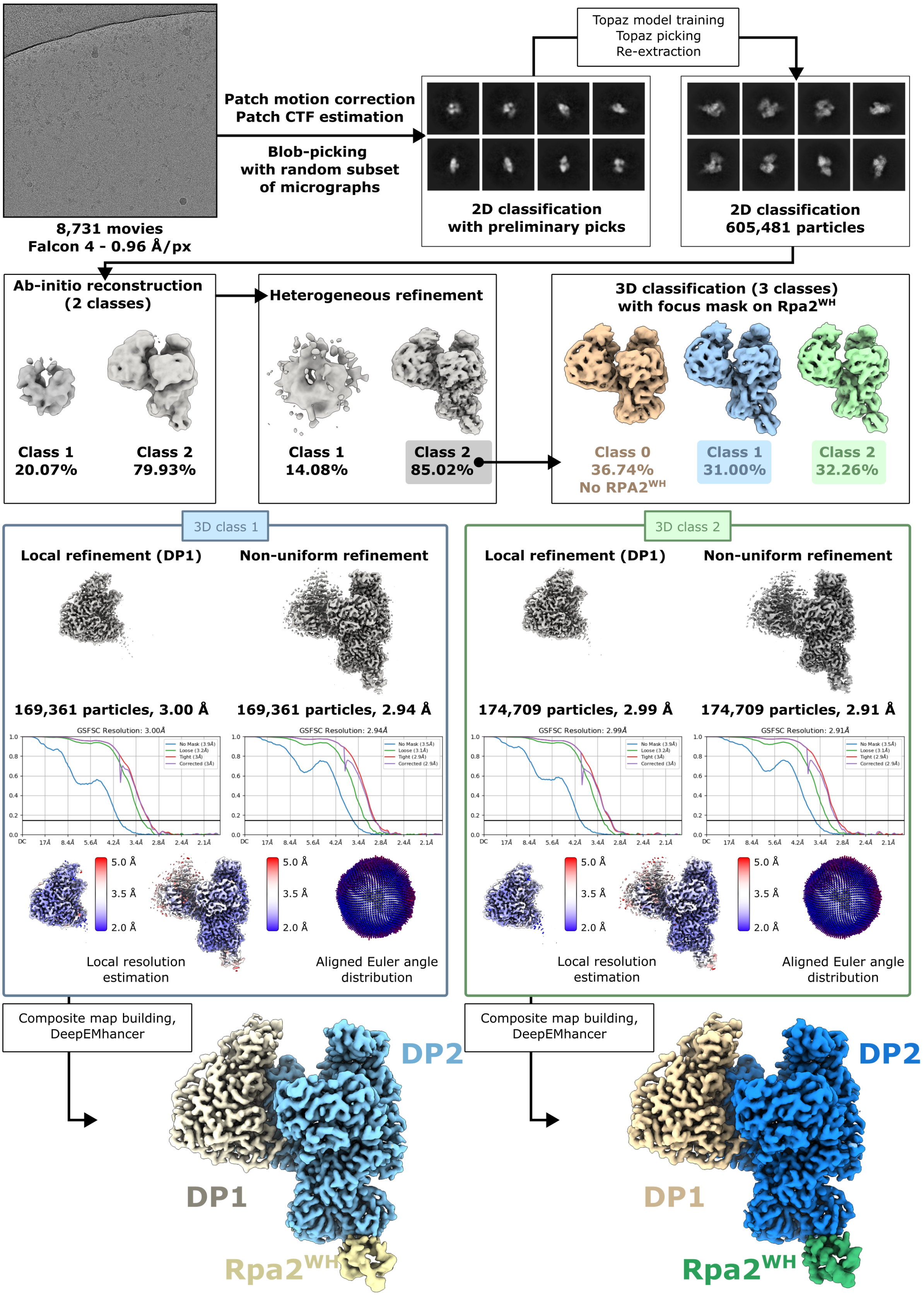

**Supplementary Figure 8**: **Cryo-EM workflow.**

| | | | |
|---|---|---|---|
| **Constraints** | | **Mean of pairwise RMSD (Å)** | |
| Dihedral restraints (ϕ) | 120 | Backbone atoms N, CA, C', O [a] | 0.42 ± 0.12 |
| Hydrogen bonds | 32 | Heavy atoms | 1.51 ± 0.14 |
| Distance restraints [a] | 954 | | |
| **Distance constraints [a,b]** | | **Energies (kcal/mol)** | |
| Unambiguous restraints | 685 | Total | -3540 ± 120 |
| Ambiguous distance restraints | 269 | Van der Waals | -342 ± 13.1 |
| Intra-residue $\lvert j-i \rvert = 0$ | 494 | Electrostatic | -3351 ± 121 |
| Sequential $\lvert j-i \rvert = 1$ | 238 | **Ensemble Ramachandran plot (% Residues)** | |
| Medium range $2 \leq \lvert j-i \rvert \leq 4$ | 129 | Most favored regions | 95.9% |
| Long range $\lvert j-i \rvert > 4$ | 93 | Additionally allowed | 4.1% |
| **Residual distance constraint violations** | | | |
| Number ≥ 0.5 Å | 10 ± 1 | **Structure Z scores** | |
| Number ≥ 0.3 Å | 12 ± 1 | Second-generation packing quality | 1.1 ± 0.2 |
| Number ≥ 0.1 Å | 18 ± 1 | Ramachandran plot appearance | 2.2 ± 0.3 |
| **RMS deviation from nOes (Å)** | 0.30 ± 0.02 | Chi1/Chi2 rotamer normality | 1.6 ± 0.8 |
| **Unsatisfied H-bond donors [c]** | 4.7 | Backbone conformation | -4.7 ± 1.5 |
| **Unsatisfied H-bond acceptors [c]** | 0 | | |

**Supplementary Table 1. Statistics for the ensemble of 10 structures calculated for full-length Rpa2$^{WH}$**. [a] For well-ordered residues (205-268). [b] For residues 175-204, only 29 intra or sequential distance constraints. [c] Per molecule. PDB ID: 9F27; BMRB ID: 34913.

| | PriSL$^{\Delta CTD}$ | PriSL$^{\Delta CTD}$+Rpa2$^{WH}$ |
|---|---|---|
| **Data collection** | | |
| Space group | *P* $4_32_12$ | *P* $2_1\ 2_12_1$ |
| Cell dimensions | | |
| *a*, *b*, *c* (Å) | 116.6, 116.6, 121.1 | 68.8, 101.5, 177.8 |
| α, β, γ (°) | 90, 90, 90 | 90, 90, 90 |
| Wavelength (Å) | 0.98011 | 0.97857 |
| Resolution (Å) | 39.5 - 1.85 (1.89 - 1.85) | 48.8 - 3.50 (3.59 - 3.50) |
| Estimated resolution limit (Å)* | 1.94, 1.94, 1.84 | 3.15, 3.59, 5.93 |
| $R_{pim}$ | 0.023 (0.609) | 0.036 (4.24) |
| $R_{merge}$ | 0.114 (4.06) | 0.126 (15.0) |
| $I / \sigma I$ | 12.6 (0.8) | 9.5 (0.3) |
| $CC_{1/2}$ | 0.999 (0.592) | 1.0 (0.20) |
| Completeness (%) | 100 (100) | 99.9 (99.5) |
| Redundancy | 26.1 (24.8) | 13.2 (13.4) |
| $R_{pim}$* | 0.025 (0.427) | 0.022 (0.506) |
| $R_{merge}$* | 0.125 (2.14) | 0.074 (1.774) |
| $I / \sigma I$* | 14.0 (1.5) | 16.1 (1.8) |
| $CC_{1/2}$* | 0.999 (0.745) | 1.000 (0.593) |
| Completeness (%)* | 96.4 (61.5) | 58.1 (16.0) |
| **Refinement** | | |
| Resolution (Å) | 39.51 - 1.85 | 48.8 - 3.50 (3.67-3.50) |
| No. reflections | 67117 (1284) | 8875 (386) |
| $R_{work}$ / $R_{free}$ (%) | 19.87/22.37 | 23.45/25.55 |
| No. atoms | | |
| Protein | 4606 | 5289 |
| Ligand/ion | 9 | 1 |
| Water | 611 | - |
| *B*-factors | | |
| Protein | 43.1 | 184.1 |
| Ligand/ion | 54.3 | 44.13 |
| Water | 57.0 | - |
| R.m.s. deviations | | |
| Bond lengths (Å) | 0.009 | 0.0085 |
| Bond angles (°) | 0.89 | 1.32 |
| PDBID | 9F28 | 9F26 |

**Supplementary Table 2. Crystallographic and refinement statistics.** Values in parentheses are for highest-resolution shell. Dataset from single crystal used per structure. Values calculated after truncation by STARANISO (*). Estimated resolution limits along the three crystallographic directions a*, b*, c*.

| Data collection and processing | **PolD-Rpa$^{WH}$ class 1**<br>EMDB-50140<br>PDB 9F29 | **PolD-Rpa$^{WH}$ class 2**<br>EMDB-50143<br>PDB 9F2A |
|---|---|---|
| Magnification | 130,000X | 130,000X |
| Voltage (kV) | 300 | 300 |
| Electron exposure ($e^-/Å^2$) | 40 | 40 |
| Pixel size (Å) | 0.96 | 0.96 |
| Symmetry imposed | - | - |
| Initial particle images (no.) | 605,481 | 605,481 |
| Final particle images (no.) | 169,361 | 174,709 |
| Map resolution (Å) (FSC threshold) | 2.94 (0.143) | 2.91(0.143) |
| Map resolution range (Å) | 2-5 | 2-5 |
| **Refinement** | | |
| Initial model used (PDB code) | 6T8H | 6T8H |
| Model resolution (Å) (FSC threshold) | 2.8 (0.5) | 2.9 (0.5) |
| Model resolution range (Å) | 2-5 | 2-5 |
| Map sharpening *B* factor ($Å^2$) | 97.2 | 96.7 |
| Model composition | | |
| Non-hydrogen atoms | 13520 | 13557 |
| Protein residues | 1688 | 1692 |
| Ligands | 4 | 4 |
| *B* factors ($Å^2$) | | |
| Protein | 58.57 | 58.79 |
| Ligand | 101.13 | 116.38 |
| RMS deviations | | |
| Bond lengths (Å) | 0.003 | 0.004 |
| Bond angles (°) | 0.488 | 0.549 |
| Validation | | |
| Molprobity score | 1.38 | 1.24 |
| Clashscore | 3.38 | 3.12 |
| Poor rotamers (%) | 0 | 0 |
| Ramachandran plot | | |
| Favored (%) | 96.3 | 97.26 |
| Allowed (%) | 3.7 | 2.74 |
| Disallowed (%) | 0 | 0 |

**Supplementary Table 3. Cryo-EM data collection, refinement and validation statistics.**

| Oligonucleotides | Length (nt) | Séquences (5' to 3') | Label |
|---|---|---|---|
| **Primer** | 17 | TGCCAAGCTTGCATGCC | 5'Cy5 |
| **Template** | 87 | CAGGAAACAGCTATGACCATGATTACGAAT TCGAGCTCGGTACCCGGGGATCCTCTAGAG TCGACCTGCAGGCATGCAAGCTTGGCA | |
| **Ladders** | 87 | TGCCAAGCTTGCATGCCTGCAGGTCGACTCT AGAGGATCCCCGGGTACCGAGCTCGAATTC GTAATCATGGTCATAGCTGTTTCCTG | 5'Cy5 |
| | 57 | TGCCAAGCTTGCATGCCTGCAGGTCGACTCT AGAGGATCCCCGGGTACCGAGCTCGA | 5'Cy5 |
| **Oligonucleotide competitor** | 87 | TGCCAAGCTTGCATGCCTGCAGGTCGACTCT AGAGGATCCCCGGGTACCGAGCTCGAATTC GTAATCATGGTCATAGCTGTTTCCTG | |

**Supplementary Table 4. Oligonucleotides used as ladders and primer-templates in the activity assay experiments.**

| Strains | Genotype or other relevant characteristics | Source or reference |
|---|---|---|
| ***E. coli*** | | |
| DH5α | *Φ80dlacZΔm15, recA1, endA1, gyrA96, thi-1, hsdR17 ($r_k$−, $m_k$+), supE44, relA1, deoR, Δ(lacZYA-argF)U169* | Thermo Fisher Scientific |
| ***T. barophilus*** | | |
| UBOCC-M-3300 | Δ*TERMP_00517* | (Birien et al., 2018) |
| **Plasmids** | | |
| pUPH | Pop-in Pop-out vector | (Birien et al., 2018) |
| pRD603 | pUPH-RPAΔ70AA Cter | This study |
| pRD605 | pUPH-RPAΔ3SU | This study |

**Supplementary Table 5: Strains and plasmids used in this study.**

| | | |
|---|---|---|
| **761-RPA32ΔWHCterKpnI** | ATCG**GGTACC**AGATCAGGCGTAGAAAGCCAGG | To construct RPA$\Delta^{WH}$ |
| **762-RPA32ΔWHCterFusRv** | CCAATTTCTTTTTTACAAAAAGAGGAATAAGAAGTCATTCTTCCTCAAATATTTCCTCCTCTAAGGC | |
| **763-RPA32ΔWHCterFusFw** | GCCTTAGAGGAGGAAATATTTGAGGAAGAATGACTTCTTATTCCTCTTTTTGTAAAAAAGAAATTGG | |
| **764-RPA32ΔWHCterBamHI** | GCTA**GGATCC**GGCTCAACATATATGCGTTTATTCTGGC | |
| **766-RPA32ΔWHVerifRv** | GGTATGCGATGCTCTTATTTGTTGTTGG | Used with 776. To verify mutant of RPA |
| **771-RPA-SupBamHI-Fw** | CCCTGCACTTATCCCCGAGAATCCATTTTCCAAGGATTATCTTCTCC | To suppress *Bam*HI site in the sequence encoding RPA |
| **772-RPA-SupBamHI-Rv** | GGAGAAGATAATCCTTGGAAAATGGATTCTCGGGGATAAGTGCAGGG | |
| **773-RPAΔ3SU-KpnI** | ATGC**GGTACC**AGTGACAGTCCCGCATGATAGG | To delete the three RPA genes |
| **774-RPAΔ3SU-FusRv** | GGTCATTTTGCAAAATCTGGAGCCTTCTTATTCCTCTTTTTGTAAAAAAGAAATTGG | |
| **775-RPAΔ3SU-FusFw** | CCAATTTCTTTTTTACAAAAAGAGGAATAAGAAGGCTCCAGATTTTGCAAAATGACC | |
| **776-RPAΔ3SUVerifFw** | GGGTTAGTATTCTAATTTTACCTCTCTTCAAAGCG | Used with 766. To verify mutant of RPA |

**Supplementary Table 6: List of primers used in this study.** Restriction sites are shown in bold.
